# Single-cell transcriptomics-guided development of flow cytometric tests predicting chronic myeloid leukemia blast crisis transformation at chronic phase diagnosis

**DOI:** 10.1101/2025.01.09.632094

**Authors:** Mengge Yu, Vaidehi Krishnan, Pavithra Shyamsundar, Kian Leong Lee, Florian Schmidt, Xi Ren, Charles Chuah, Shyam Prabhakar, Verity Saunders, David Yeung, Naranie Shanmuganathan, Agnes S.M. Yong, Timothy P. Hughes, S. Tiong Ong

## Abstract

Clinical risk scores in chronic myeloid leukemia (CML) are inadequate for identifying chronic phase (CP) patients at high-risk of blast crisis (BC) progression. The lack of accurate predictive tests hamper timely interventions, including stem cell transplants, that are more effective in early disease. By interrogating a single cell atlas of primary imatinib resistance for a BC-like gene expression signature, we identify aberrant CD42A^+^ megakaryocytic and CD10^+^CD19^+^ lymphoid progenitor expansion, as well as STAT1- and IFNγ-related inflammatory programs, as consistent features in the bone marrow and peripheral blood of CP patients at high-risk of BC transformation. We develop multi-color flow cytometry-based tests (MFC) to detect these features in CP patients at the time of diagnosis. Validating our MFC panels on a combined Australia-Singapore cohort comprising 28 CP patients, including 14 who underwent transformation, we demonstrate their ability to detect 100% of CP patients who transform with no false positives. Our findings highlight small populations of inflamed hematopoietic stem and progenitor cells, present at CP diagnosis, as powerful harbingers of future BC transformation. The ability of MFC panels to detect these cells at diagnosis support their inclusion as accurate risk-assessment tools to improve management of high-risk patients.

## Introduction

Blast crisis (BC) chronic myeloid leukemia (CML) remains a major challenge in the management of CML patients as survival is usually measured in months. The ability to confidently identify chronic phase (CP) patients at high-risk of BC transformation may be life-saving, since it would prompt closer clinical monitoring and more aggressive treatment during CP, including stem cell transplantation, when such modalities are more effective.[1, 2] However, high-confidence predictions for future BC transformation are not currently possible, and will require the elucidation of pre-treatment factors that determine disease progression. Toward this goal, we had previously identified a convergent BC transcriptome that is shared between lymphoid and myeloid BC CD34^+^ hematopoietic stem and progenitor cells (HSPC),[3] and which is also detectable in HSPCs from CP patients at increased risk of developing TKI resistance as well as earlier BC transformation.[4-6] More recently, we published a single-cell atlas of primary TKI resistance (which we term the ‘Resistance Atlas’) that identified three cellular Hallmarks predicting BC transformation at CP diagnosis: lineage skewing away from TKI-sensitive erythroid progenitors and toward lymphoid or myeloid progenitors, an inflammatory gene expression (GE) signature among progenitors, and a reduction in adaptive NK cells.[4] Here, we further interrogate the Resistance Atlas to identify cell-based markers of these Hallmarks detectable by multi-color flow cytometry (MFC) (**Supplementary Table 1**). Thereafter, using an independent cohort of CP patients, we determine the clinical validity of the assembled MFC antibody panels to segregate patients destined to undergo BC transformation (CP^BC^) from those with optimal tyrosine kinase inhibitor (TKI) responses (CP^Resp^) using diagnostic BM and/or PB samples (**Figure 1A** and **Supplementary Table 2**).

**Figure 1.**
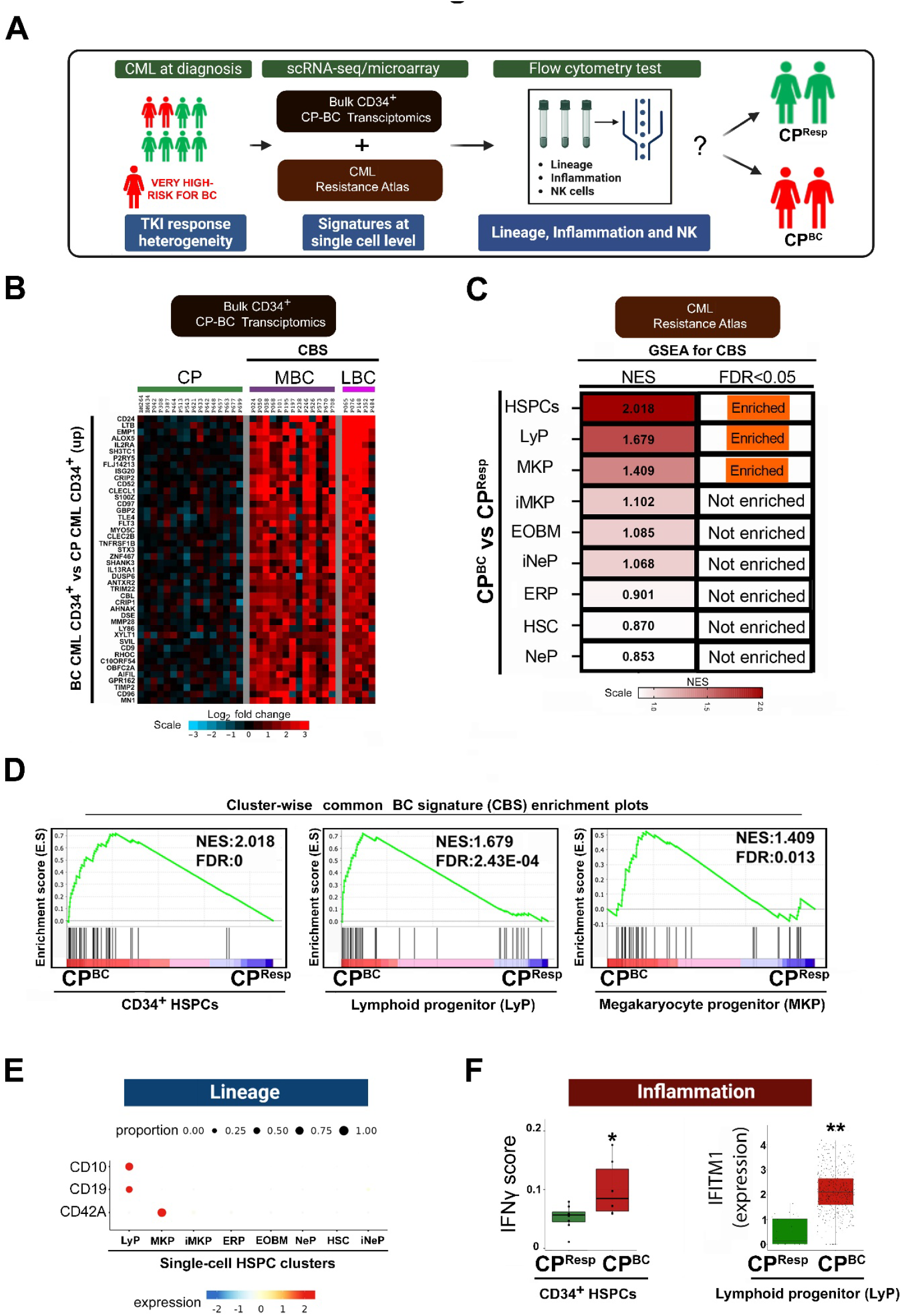
Single cell transcriptomic-based design of flow cytometric panels to detect high-risk CP patients. **A**. Study Design: Development of a flow cytometry-based clinical test to categorise CML patients at the time of diagnosis into 2 prognostic categories: patients with optimal European LeukemiaNet[19] TKI responses (CP^Resp)^, and those who eventually transformed to BC (CP^BC^). The flow cytometry test would comprise of three main features of TKI resistance i.e. lineage skewing away from erythroid lineage, inflammation and adaptive NK cell accumulation defect, and our study tests their predictive power. **B**. Derivation of a common BC gene expression signature (CBS). Genes which were commonly upregulated in CD34^+^ myeloid (MBC) and lymphoid (LBC) samples compared to CD34^+^ CP samples (>2.5 log_2_fold change, p-adjust≤0.05) were derived from a previously published microarray-based dataset[3], and used to define the CBS. **C**. Identification of cell clusters enriched for the CBS within the single cell Resistance Atlas. GSEA for the CBS was performed comparing different cell clusters from CP^BC^ and CP^Resp^ samples, and the normalized enrichment scores (NES) plotted as a heatmap. Comparisons with FDR q-value ≤0.05 were considered significantly enriched. LyP, lymphoid progenitor; MKP, megakaryocyte progenitor; iMKP, immature megakaryocyte progenitor; EOBM, eosinophil-basophil mast cell progenitor; iNeP, immature neutrophil progenitor; Pro-B cells, Pro-B; ERP, erythroid progenitor; HSC, hematopoietic stem cell; NeP, neutrophil progenitor. **D**. CBS enrichment plots for HSPCs, LyPs, and MKPs from CP^BC^ and CP^Resp^ samples from (**C**). **E**. Individual CD34^+^ cell clusters were subjected to marker gene analysis using the Seurat pipeline, and cell surface markers used as lineage read-outs in our MFC panels. The bubble plot shows expression (color scale) while the size of the bubble represents the percent positivity of the indicated genes within individual cell clusters. **F**. (Left) Module scores for the ‘Hallmark IFNγ’ gene set was computed for pseudobulked aggregated CD34^+^ HSPCs from CP^BC^ and CP^Resp^ samples. Module score p-values were determined by the Kruskal Wallis test (*p*<0.05). (Right) Box plot shows the expression of the *IFITM1* gene within LyPs from CP^BC^ and CP^Resp^ samples. Gene expression p-values were compared by the Wald test (*p*<0.01).

## Methods

### Patient cohort, sample preparation, flow cytometry, and data analysis

CML patient samples were collected following Institutional Review Board (IRB) approval and patient consent. The cohort in this study comprises 14 patients each in the CP^Resp^ and CP^BC^ groups, with the latter including 7 each transforming to myeloid and lymphoid BC with a duration spanning 3-91 months (**Supplementary Table 2**). Diagnostic BM or PB derived mononuclear cells (MNCs) samples were stained for flow cytometry using specific antibody panels that detected lineage (Panels 1 and 2), inflammation (Panel 3 and 4) and adaptive NK cells (Panel 5). All samples were analysed on the BD LSR Fortessa™ cell analyzer (BD Biosciences). MFC data were analyzed using the FlowJo software, and significance was assessed using the 2-tailed Student’s t-test.

## Results and Discussion

We interrogated the CML Resistance Atlas to determine which sub-clusters within the HSPC compartment of CP^BC^ patients harbored the BC-like transcriptome we had previously identified.[3] First, by comparing bulk CD34^+^ CP and BC transcriptomes,[3] we derived a novel gene set comprising 44 of the most differentially upregulated genes in both myeloid and lymphoid BC HSPCs compared to CP HSPCs, which we term the ‘common BC signature’ (CBS) (**Supplementary Table 3** and **Figure 1B**). As expected, pseudobulked HSPCs from CP^BC^ samples within the Resistance Atlas exhibited significant enrichment of the CBS compared to CP^Resp^ samples (**Figure 1C**). Next, focussing on individual populations within the HSPC compartment of CP^BC^ samples, we identified megakaryocytic (MKP) and lymphoid (LyP) progenitor clusters, to be enriched for the CBS (**Supplementary Figure 1A, Figure 1C** and **1D**). These findings prompted us to include markers for MKP and LyP populations as indicators of prognostic sub-populations expanding in TKI-resistant CML. Using inter-cluster marker gene analysis, we identified *GP9* (CD42A) as an MKP marker, and *MME* (CD10) and *CD19* as LyP markers (**Figure 1E**), and incorporated these into panels designed to detect MKP (Panel 1) and LyP (Panel 2) expansion. For Inflammation panels, we had previously noted that CP^BC^ HSPCs display a resistance-conferring inflammatory state characterized by activation of the IFNγ pathway and its downstream effector STAT1 in CP^BC^ patients who transform to either MBC or LBC (**Figure 1F, left**).[4, 7] Because STAT1 phosphorylation at serine-727 is a readout for STAT1 activity, we included an antibody directed against pSTAT1(Ser-727) within the HSPC population (Panel 3).[8] To identify additional markers of inflammation, we conducted differential gene expression analysis on HSPC clusters that harbour the CBS. Here we found IFITM1 (CD225), an IFNγ-induced inflammatory marker,[9] to be specifically upregulated within LyP cells (**Supplementary Table 4**), and created a LBC-specific Inflammation panel (Panel 4) (**Figure 1F, right**). Lastly, based on previous literature, we designed a flow cytometry panel as a read-out for adaptive NK cell abundance, specifically incorporating the activating NK cell receptor, NKG2C (Panel 5).[10]

Next, we assessed the ability of the MFC panels, individually or in combination, to segregate CP^BC^ from CP^Resp^ in an independent cohort of 28 CP patients from the TIDEL II study[11] (n=23) and Singapore (n=5) (**Supplementary Figure 1B** and **Supplementary Table 5**). The gating strategies for the 5 MFC panels and populations being assayed are displayed in **Supplementary Figures 2** and **3**. Optimal cut-off points maximizing specificity and sensitivity for each biomarker panel were determined by calculating the Youden’s J index along the ROC curve (**Supplementary Table 6**),[12] which was then used to classify patients as being positive or negative for each panel (**Online Methods**). Using Panel 1, which measures CD42A^+^ MKPs, we correctly classified 75% (9/12) of CP^BC^ patients (CP^MBC^ *p*=0.04, CP^LBC^ *p*=0.01;), with no false positives among CP^RESP^ patients (**Figure 2A, left**). With Panel 2, we detected CD10^+^CD19^+^ LyP expansion in 6/7 (85.7%) CP^LBC^ patients, with no false positives among CP^RESP^ or CP^MBC^ patients (**Figure 2A, right**). Together, Panels 1 and 2 demonstrate aberrant lineage expansion as a sensitive (78.6%, 11/14) and specific (0% false positive) marker of CP^BC,^ but with four outlier samples, S15, S16, S25 and S28, which could not be identified based on lineage Panels 1 and 2 alone. Next, we evaluated the ability of Panels 3 and 4 to detect progenitor inflammation among CP^BC^ patients, and potentially, improve on lineage expansion-based identification. Using Panel 3, which assesses pSTAT1(Ser-727), we observed increased HSPC inflammation in all CP^MBC^ patients (7/7) and the majority of CP^LBC^ patients (5/7), but none among 14 CP^RESP^ samples (*p*<0.01 for CP^MBC^; *p*=0.07 for CP^LBC^; **Figure 2B, left**). Using Panel 4, which detects LyP-specific inflammation through IFITM1, we found that all CP^LBC^ (7/7) samples were positive while no healthy controls were positive (*p*<0.01; **Figure 2B, right**). Together, Panels 3 and 4 detected 100% (14/14) of CP^BC^ patients with no false positives among CP^RESP^ patients, and importantly correctly predicted the outlier samples from Panels 1 and 2, S15, S16, S25 and S28 as being destined for BC. Lastly, using Adaptive NK Panel 5, we confirmed that NKG2C^+^ adaptive NK cells were significantly higher in CP^RESP^ compared to CP^BC^ patients (**Figure 2C**) (*p*<0.05).

**Figure 2.**
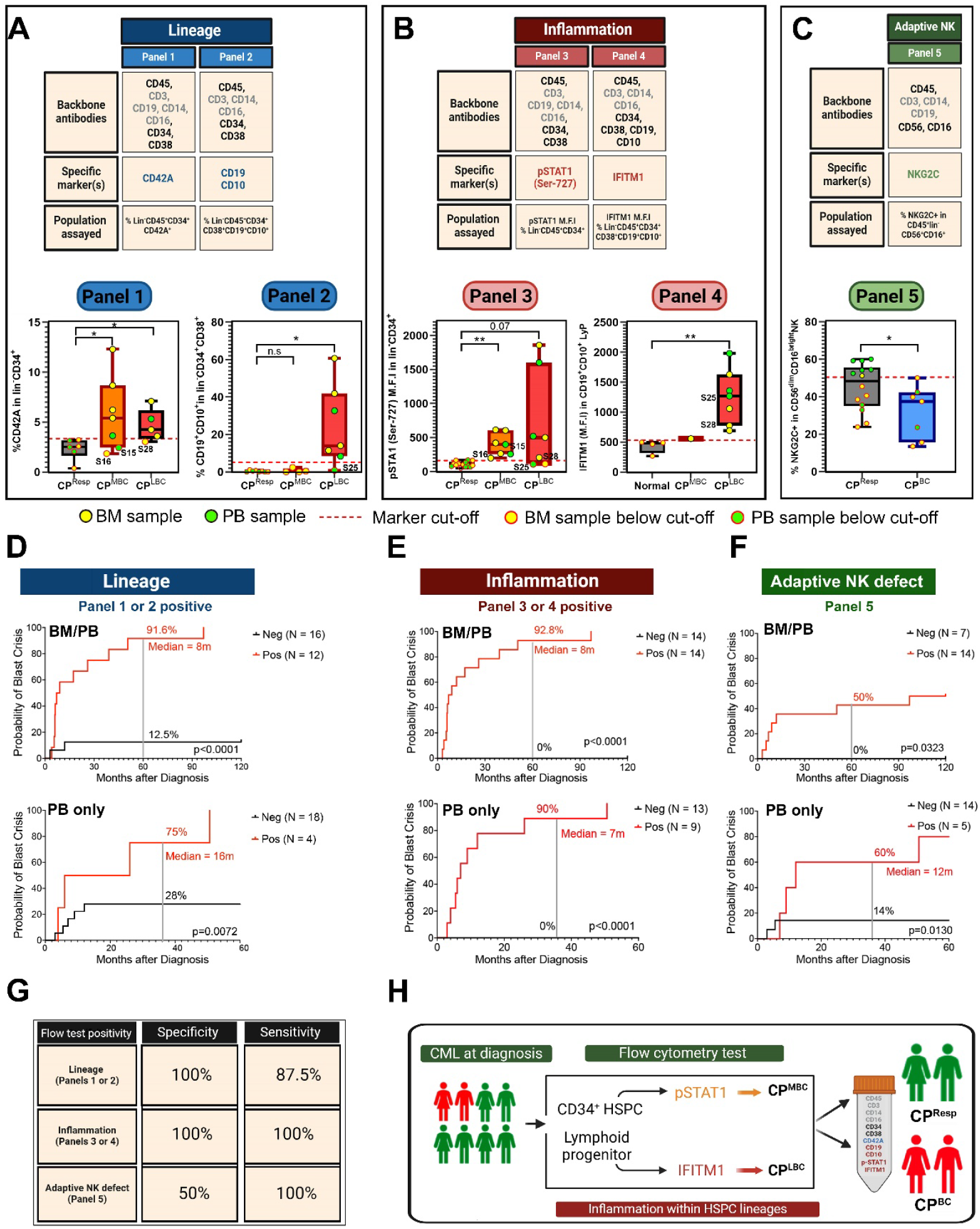
Inflammation within HSPCs is a powerful predictor of BC transformation at the time of CP diagnosis. **A-C**. BM (yellow symbols) or PB MNCs (green symbols) from a cohort of 28 CP CML patients who were either optimal TKI responders (CP^Resp^) or who underwent MBC (CP^MBC^) or LBC (CP^LBC^) transformation were subjected to flow cytometry using the depicted antibody panels. Red dotted line depicts the Youden’s J index cut-off values calculated for each panel (see **Online Methods**). **A**. Flow cytometry panel for detection of lin^-^CD34^+^CD42A^+^ MKPs (Panel 1) and lin^-^ CD34^+^CD38^+^CD19^+^CD10^+^ LyPs (Panel 2). Box plots show the percentages of the indicated populations. **B**. Flow cytometry panels for detection of HSPC (Panel 3) or LyP (Panel 4) inflammation. Left: Mean fluorescence intensity (M.F.I) of pSTAT1 (Ser-727) within lin-CD34^+^ population. Right: M.F.I of *IFITM1* within the LyP identified as lin^-^CD34^+^CD38^+^CD19^+^CD10^+^. Note: For M.F.I calculations, populations<20 cells were excluded. Since all CP^Resp^ samples (n=8) had less than 20 LyPs, these data could not be plotted, and LyPs from normal BM (normal) was plotted as a comparison. **C**. Flow cytometry panel (Panel 5) for detection of adaptive NK cells. NKG2C^+^ adaptive NK cell abundance within CD56^dim^CD16^bright^ NK cells were plotted. Two-tailed Student’s t-test was used for all comparisons. **D-F**. Kaplan-Meier curves were plotted using positive or negative calls as determined by the Youden’s J index (**Supplementary Table 7** for complete data), and based on lineage Panels 1 and 2 (**D**), inflammation Panels 3 and 4 (**E**), or the adaptive NK Panel 5 (**F**). The y-axis indicates the probability of BC transformation, while the x-axis the time taken to transformation. Kaplan-Meier curves were compared using the Log-rank (Mantel-Cox) test. **Top**: All samples including BM and PB were used for the plots. Note: When both BM and PB samples were available (n=7), only the BM sample was chosen for the top plots. p<0.001 for **D** and **E** and p=0.03 for **F. Bottom**: PB samples were used to generate Kaplan-Meier curves. p<0.001 for **D** and **E** and p=0.0130 for **F. G**. The sensitivity and specificity values for the MFC panels detecting at-risk lineage, inflammation or adaptive NK cell features are tabulated (using BM and PB data). **H**. A flow cytometry-based clinical test that can quantify inflammation within HSPC lineages emerges as a powerful predictor of BC transformation at the time of CP diagnosis.

Next, we binarized the MFC data, based on marker positivity cut-offs as positive or negative for the three Hallmark features, and generated Kaplan-Meier curves to assess the performance of the five MFC panels (**Figure 2D and Supplementary Table 7**). When CP patients were segregated according to Lineage Panels 1 and 2, the specificity and sensitivity for predicting BC transformation was 100% and 90%, respectively (*p*<0.0001 by log-rank test) (**Figures 2D, top)**. Strikingly, when using Inflammation

Panels 3 and 4, the specificity and sensitivity were both 100% with 92.8% of positive patients at the time of diagnosis likely to transform within 5 years (*p*<0.0001 by log-rank test) (**Figure 2E, top**). While segregation of diagnostic samples based on NK Panel 5 had a specificity of 100%, sensitivity was only 50% (p<0.03 by log-rank test) (**Figure 2F, top**). Consistent results were obtained when flow panel performances were evaluated using only diagnostic PB MNCs (**Figure 2D-F**, bottom plots), with 90% of inflammation positive patients at the time of diagnosis likely to transform within 5 years (*p*<0.0001 by log-rank test) (**Figure 2E, bottom**).

In summary, our results are clinically important because they demonstrate the ability, using standard MFC, to identify patients at high-risk of BC transformation at diagnosis with high specificity and sensitivity (**Figure 2G**). Based on our combined results, we propose the use of a primary 10-antibody tube (integrating Panels 2-4) for assessing inflammation within the HSPC and LyP compartments (**Figure 2H**). The detection of both lineage skewing as well as HSPC inflammation are important to achieve high sensitivity for detecting CP^BC^ patients, and together, may be regarded as prognostic ‘red flags’ that prompt more intensive management strategies. Accordingly, Panel 3 detected inflamed HSPCs even when MKP expansion was not evident, including two CP^MBC^ patients (S15, S16) and one CP^LBC^ patient (S28), while Panel 4 identified inflammation in the patient S25 who did not show LyP expansion. Here, we note that two CP^LBC^ samples were pSTAT1 negative, suggesting that IFITM1 is a more sensitive marker for inflammation in CP^LBC^ patients. Importantly, our MFC panels also identified 3 CP^BC^ patients with low clinical risk scores, all of whom transformed within 7 months of diagnosis, suggesting their utility in detecting patients who undergo ‘sudden-onset BC’[13] (**Supplementary Table 2**). Our results are also biologically informative, and implicate activation of inflammatory pathways, including INFγ-STAT1, as early progression events preceding progenitor expansion. Analogous to the CBS, inflammation thus represents a biological convergence point shared across high-risk HSPC populations, suggesting an important role in drug resistance and disease progression to both MBC and LBC. Our finding of MKP expansion in CP^BC^ patients is also consistent with reports implicating increased BM megakaryocytes and high platelet counts as poor prognostic factors.[14, 15] Future studies are warranted to test the clinical utility of our MFC panels in larger prospective cohorts, to better understand the causes and consequences of inflammation in CML, and also to integrate MFC-based tests with genomic tests to predict CML treatment outcome.[16-18]

## Acknowledgements

We are grateful for the assistance of Dr. Roger Vaughn, Duke-NUS Medical School, in performing the statistical analyses in Figures 2D-F. Figures were created with Biorender.com.

## Authorship Contributions

Contribution: M.Y. contributed to flow cytometry panel design, data acquisition, data analysis and interpretation. V.K. contributed to study design, data analysis and interpretation, and manuscript writing; P.S and K.L.L contributed to data analysis and interpretation during MFC panel design; K.L.L contributed to data analysis and interpretation during MFC panel design; F.S. and R.X. contributed towards scRNA-seq gene expression analysis; S.P. contributed to data interpretation and analysis for the scRNA-seq project; C.C., V.S., D.Y., N.S., A.S.M.Y., and T.H. provided annotated clinical samples and participated in discussions; S.T.O. conceived and oversaw the project, while also providing financial support, data interpretation, and manuscript writing.

